# Impact of Grain for Green Project on Water Resources and Ecological Water Stress in Yanhe River Basin

**DOI:** 10.1101/2021.10.25.465705

**Authors:** Yuping Han, Fan Xia, Huiping Huang, Wenbin Mu

## Abstract

Grain for Green project (GGP) initialed by China government since 1999 has achieved substantial achievements accompanied with surface runoff decrease in the Loess Plateau but impacts of large-scale afforestation on regional water resources are uncertain. Hence, the objective of this study is to explore the impact of land use change on generalized water resources and ecological water stress using blue and green water concept taking Yanhe River Basin as a case study. Soil and Water Assessment Tool (SWAT) is applied to quantify summary of green and blue water which is defined as generalized water resources, ecological water requirement of vegetation (forest and grass), agricultural water footprint and virtual water flow are considered as regional water requirements. Land use types of 1980 (scenario I), 2017 (scenario II) are input in SWAT model while keeps other parameters constant in order to isolate the influence of land use changes. Results show that average annual difference of blue, green and generalized water resources is −72.08 million m^3^, 24.34 million m^3^, −47.74 million m^3^ respectively when simulation results of scenario II subtracts scenario I and it presents that land use change caused by GGP leads to decrease in blue and generalized water resources whereas increase in green water resources. SURQ in scenario I is more than that in scenario IIin all the study period from 1980-2017, green water storage in scenario I is more than that in scenario II in all the study period except in 1998; whereas LATQ in scenario I is less than that in scenario II except in 2000 and 2015, GWQ in 1992, 2000 and 2015, green water flow in 1998. Blue water, green water storage and green water flow in scenario II is less than that in scenario I in the whole basin, 12.89 percent of the basin and 99.21 percent of the basin respectively. Total WF increases from 1995 to 2010 because forest WF increases significantly in this period though agricultural WF and grass WF decreases. Ecological water stress index has no obvious temporal change trend in both land use scenarios but ecological water stress index in scenario II is more than that in scenario I which illustrates that GGP leads to increase of ecological water stress from perspective of generalized water resources

## Introduction

China government launched Grain for Green project (GGP) implements in the 1990s in order to control soil erosion and water loss in the Loess Plateau [1–3]. After more than two decades of vegetation restoration, soil erosion caused by unreasonable land use has been curbed, and the ecological environments of the region have been greatly improved [4–5]. Theoretically, vegetation restoration can enhance vegetation coverage, increase precipitation interception and water retention, decrease soil erosion and thus improve ecosystem stability. Forests consume more water than other vegetation types such as agricultural crops and natural grasslands [6], therefore, accompanying with the enhancement of vegetation coverage since 2000 in China, runoff shows significant decrease in Haihe, Liaohe, Songhua Jiang, Hanjiang and the Yellow River [7,8,9]. Particularly, runoff in middle reaches of the Yellow River which has the most obvious vegetation restoration achievements has reduced sharply, the value in Huayuankou station has decreased from 55.9 billion m^3^ (1970s) to 45.2 billion m^3^ (2010-2015) [10]. Some results indicate that large-scale vegetation restoration in the Loess Plateau has positive impacts on soil erosion control, ecological environment and negative impacts on streamflow [11–13]. Studies in other areas of the world also demonstrate that increase of vegetation coverage will lead to higher interception loss which is the main reason for streamflow reduction [14–15]. Some studies also indicate that unsuitable vegetation type, overlooking the bio-diversity will bring about the soil desiccation in the Loess Plateau [16–17]. There are also studies present the negative effects of afforestation on the underground water resources [18] and ecological water deficit because of afforestation [19]. Why does stream decrease and where does it go? Whether the soil turns dry and water stress becomes serious as a result of GGP? These are important problems need to be discussed.

Blue and green water concepts proposed by Falkenmark [20] provide new theories and ideas for water resources management, especially in arid and semi-arid regions. Large amount of blue water converting to green water is one of the important reasons for streamflow decline in the Loess Plateau [21]. Vegetation coverage improvement makes streamflow reduction significantly and vegetation restoration is close to the threshold of water resources carrying capacity from point view of blue water [22]. However, it can cut down water in sediment transport and virtual water embodied in green plants is far greater than streamflow reduction from perspective green water [23]. Water footprint (WF) proposed by Hoekstra and Hung [24] represents direct and indirect measurements of water appropriation by human beings. It quantifies blue and green water consumption in a river basin or a specified region and is a new approach to assess sustainable water use of economic production sectors or regions [25–27].

Yanhe River Basin in the Loess Plateau is the first tributary of the Yellow River and it is in semi-arid area with serious water scarcity and severe soil erosion. The watershed is one of the earliest and fastest areas in the whole country to reclaim cultivated land to forests (grasslands) and the vegetation restoration effect is significant since implement of GGP. During the period of 2000-2017, forests increased by 2357.6 km^2^, cultivated land decreased by 2116.2 km^2^, urban land increased by 222.1 km^2^, waters, grassland and other land cover types did not change remarkably. Numerous studies have attempted to evaluate vegetation restoration impact on water resources and focus on improvement of watershed ecological environment, amelioration of soil properties, change of streamflow and sediment [28–30]. However, there is almost no study exploring the impact of vegetation restoration on water stress from the aspect of water footprint. Therefore, the objectives of this paper is to: (1) analyze spatial-temporal characteristics of land use change during period of 1980-2017 in Yanhe River Basin and describe GGP achievements; (2) quantify water balance elements and analyze spatial-temporal characteristics of green and blue water in the whole basin and sub-basins based on calibrated and validated SWAT model simulation results; (3) investigate temporal characteristics of agricultural WF, ecological WF and virtual water flow; (4) calculate ecological water stress index based on generalized water resources and water footprint, probe impact of GGP on regional water stress.

## Materials and methods

### Study area

Yanhe River Basin (36.21’-37.19’N, 108.38’-110.29’E) is located in the Loess Plateau in northern Shaanxi province in China and is a first-order tributary of the Yellow River (Figure 1). With an area of 7785 km^2^, the watershed has warm temperate continental semi-arid monsoon climate. Annual mean precipitation of the watershed is 520 mm with 75 percent being concentrated from June to September, and the mean annual temperature varies from 8.8 to 10.2 degrees Celsius [31]. Yearly streamflow of Ganguyi is 220 million m^3^, major land use and land cover types of the watershed are forest, shrub land, grassland, cultivated land, construction land, water body, and bare land. The water resources per capita in the basin is 375 m^3^, accounting for 28 percent of Shaanxi Province and 17 percent of China. This river is with large sediment content, serious point and non-point source pollution. The water resources in the watershed are in acute shortage and the ecological environment is fragile.

**Figure 1.**
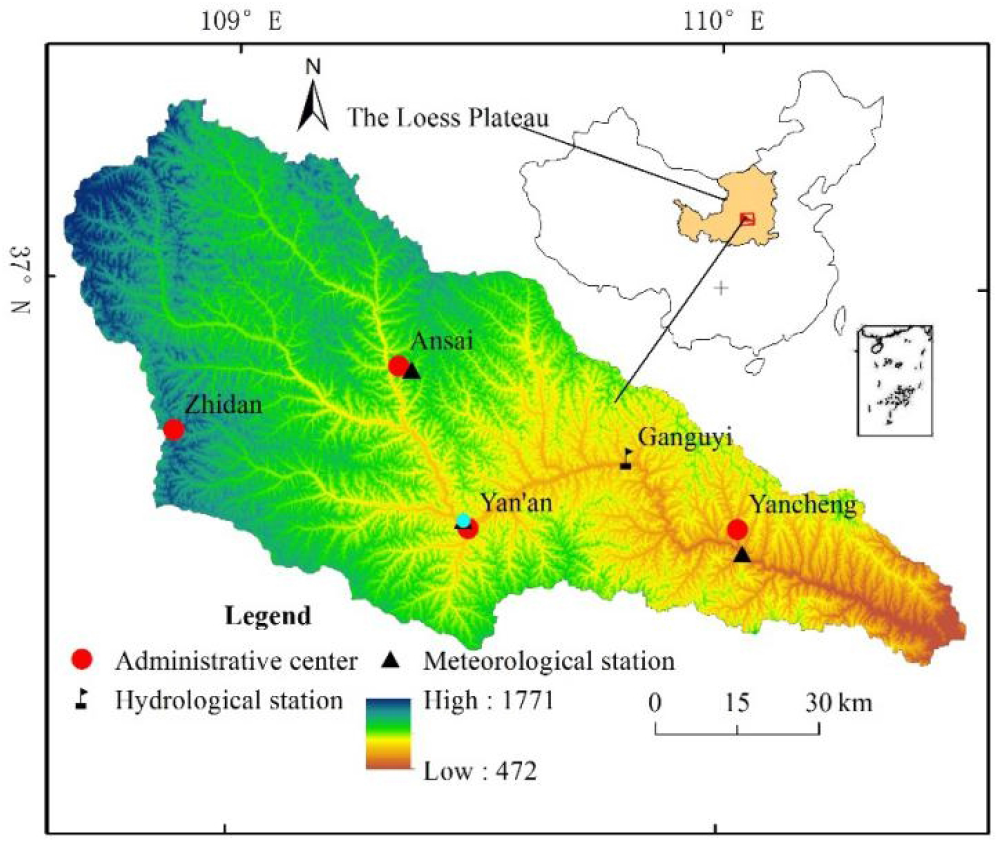
Location and distribution of hydrological and meteorological stations in Yanhe River Basin

### Data sources

Land use of the region in 1980 (before GGP) is from Resource and Environment Science and Data Center (http://www.resdc.cn), data of 2017 (after GGP) is interpretated from Landsat OLI images. The image data with 30 m resolution and cloud cover 0 are collected from Geospatial Data Cloud (http://www.gscloud.cn/). Digital evaluation model (DEM) data with 30 m resolution is also provided by Geospatial Data Cloud (http://www.gscloud.cn/). The daily meteorological data were extracted from China’s meteorological data sharing service system (http://cdc.cma.gov.cn/home.do). Soil data with an accuracy of 1:1000000 is obtained from the data center for resources and environment sciences. Annual streamflow data of hydrological station is from the Yellow River conservancy commission (YRCC). Various crop yields, population, grain consumption are obtained from Yan’an Statistical Yearbook; forest and grassland area are calculated from land use map.

### SWAT model

The Soil and Water Assessment Tool (SWAT) is a physically based, semi-distributed model. It has the advantage to simulate quality and quantity of both surface and ground water as well as to predict impact of land cover change, land management practices and climate change [32–34]. The model has been successfully applied to calculate water yield, evaluate water quality to small watershed as well as to large river basin [35–37]. Simulation results can be applied to quantify spatial and temporal characteristics of blue and green water in different parts of the world, for example Savannah River Basin in south-east Atlantic region of USA [38], Weihe River in northwest China [39], Athabasca River Basin in Canada [40].

SWAT Calibration and Uncertainty Programs (SWAT-CUP) is applied to calibrate and validate SWAT model [41]. T-stat and p-value calculated by Sequential Uncertainty Fitting programme algorithm (SUFI-2) in SWAT-CUP are used to estimate the sensibility of every parameter. T-stat represents sensitivity degree, and the greater the absolute value is, the more sensitive it is; p-value represents parameter sensitivity significance, and the closer it is to 0, the more significant it is. R^2^ (coefficient of determination) and NSE (Nash-Sutcliffe Efficiency) are used to evaluate accuracy of SWAT model. R^2^ is from 0 to 1 and if the value is close to 1, it indicates that the simulated data is close to the observed data and vice versa [42].

### Estimating blue and green water resources

Blue water resources in each sub-basin is equal to water yield (WYLD) which is the sum of surface runoff (SURQ), lateral flows (LATQ) and ground water recharge (GWQ) [43]. Green water resources consist of green water flow (actual evapotranspiration, ET) and green water storage (soil water content, SW). The annual SURQ, LATQ, GWQ, ET and SW are obtained from simulation results using the calibrated and validated SWAT model [44–45]

To investigate the influences of land use change caused by GGP on blue, green water and every hydrological element, two scenarios are set up by changing land use types while keeping other parameters input in SWAT model unchanged (scenario I : land use of 1980; scenario II: land use of 2017). Only impact of land use change is considered when simulation results in scenario II subtract data in scenario I.

### Water footprint

Water footprint (WF) concept quantifying the volumetric freshwater consumption of products is distinguished as green, blue and grey WF [20]. Green WF (WFgreen) is defined as rainwater that is stored in soil and evaporated or consumed during production. Blue WF (WFblue) refers to surface and groundwater that are consumed or evaporated during production. Grey water footprint (WFgrey) expresses the volume of water needed to dilute pollutants to achieve allowed values in the receiving water bodies. The summary of WFgreen and WFblue demonstrates quantity of total freshwater consumption during production, while WFgrey indicates degradation of water quality. The goal of this study is to explore water quantity change caused by vegetation restoration and so WFgrey is not considered during calculation process.

### Agricultural crop WF

Crop WF is the sum of WF_green_ and WF_blue_ which is determined by following equations:

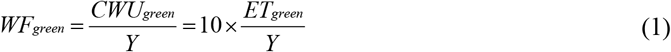

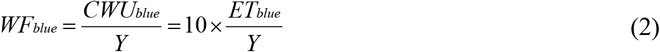

Where *WF*_*green*_, *WF*_*blu*e_ is green WF (m^3^/t) and blue WF (m^3^/t) during crop growth season respectively; *CWU*_*green*_ and *CWU*_*blue*_ are green and blue water use (m^3^/ha); 10 is a constant to convert water depth (mm) to water volume (m^3^/ha); *Y* is crop yield (m^3^/ha); *ET*_*green*_ and *ET*_*blue*_ are defined as the evaporative demand satisfied by green and blue water, respectively. *ET*_*green*_ and *ET*_*blue*_ are calculated as [46]:

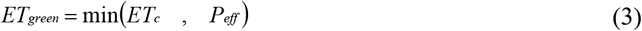

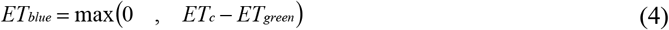

Where *ET*_*c*_ represents actual evapotranspiration of crops from sowing day to harvest; *P*_*eff*_ is effective precipitation.

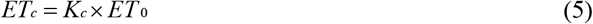

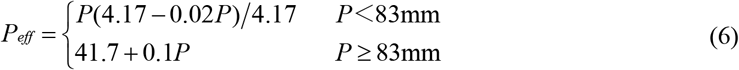

Where *K*_*c*_ is crop coefficient, *ET*_*0*_ is potential reference crop evapotranspiration and is calculated by Penman-Monteith formula; *P* is precipitation of ten days.

### Vegetation WF

The equation of vegetation WF is:

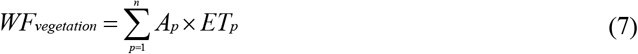

Where *WF*_*green*_ is the vegetation WF (m^3^); *A*_*p*_ is the area of vegetation coverage (m^2^); *ET*_*p*_ is the vegetation evapotranspiration (mm/day) under restricted circumstances, *p* is vegetation type.

Vegetation evapotranspiration is less than potential evapotranspiration when soil water content is below a specified threshold, and the effect is determined by the soil moisture limitation coefficient (*K*_*s*_). Thus, the calculation equation of *ET*_*p*_ is as follows:

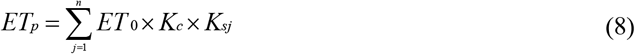

Where *ET*_*0*_ is potential reference evapotranspiration, *K*_*c*_ is the vegetation water demand coefficient, *K*_*s*_ is the soil moisture limitation coefficient, *j* is the soil type.

Based on previous studies on the Loess Plateau [47–48], *K*_*c*_ of forest and is grassland is 0.765 and 0.65 respectively. Soil type in Yanhe River Basin contains silty soil and sandy loam, *K*_*s*_ of the two soil types is 0.537 and 0.556 respectively.

### Ecological water stress index (EWSI)

Raskin [49] proposed a criterion using the ratio between water demand and available water resources to estimate water scarcity, which has been widely used to evaluate global and regional water resources [50–51]. Water stress index (WSI) can provide information on management of fresh water resources [52]. In this paper, EWSI is calculated as the ratio of ecological water footprint to generalized water resources (Figure 2). Generalized water resources is the sum of blue water and green water simulated by SWAT model, ecological water footprint contains agricultural WF, forest WF, grass WF and virtual water flow.

**Figure 2.**
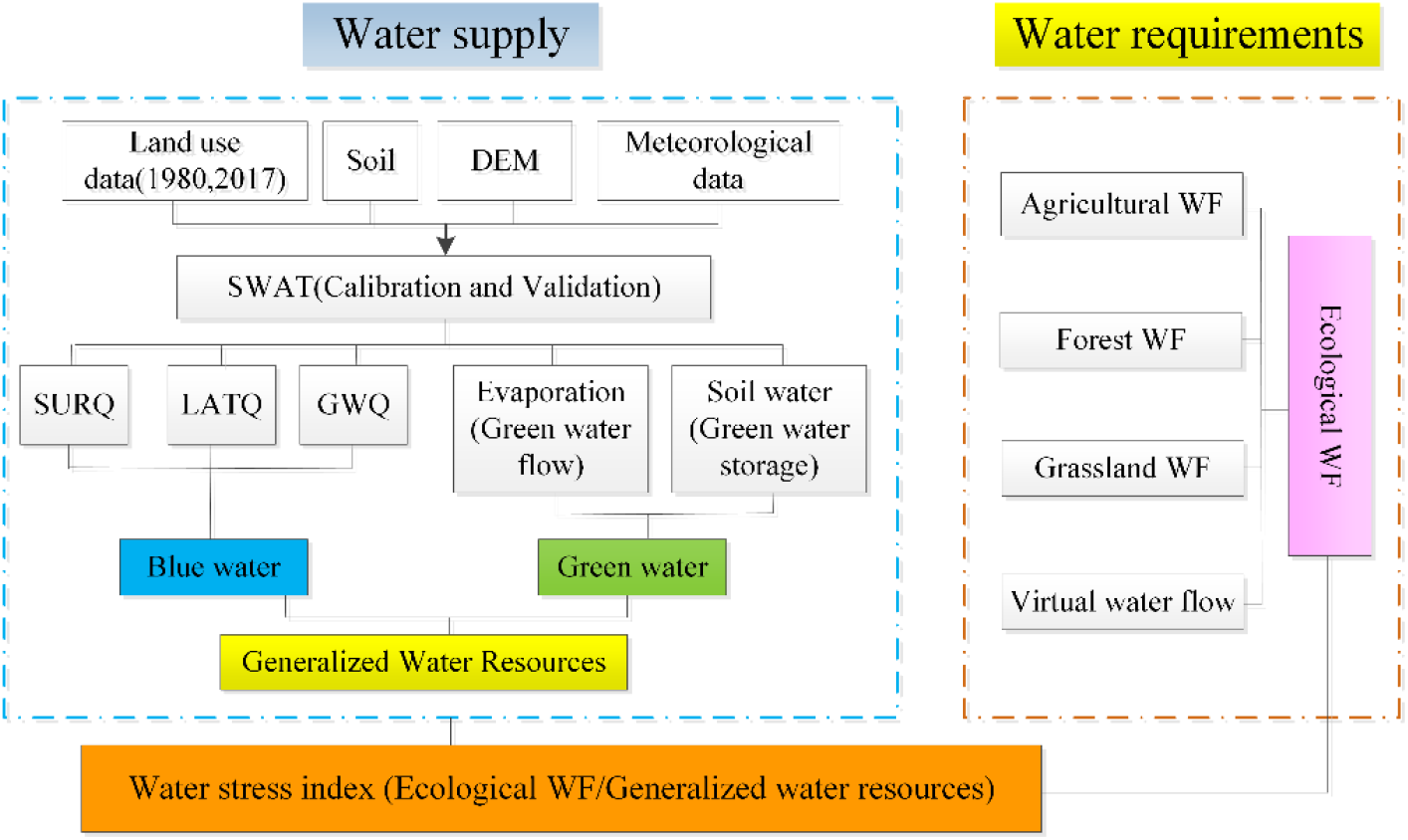
Framework for assessing ecological water stress in Yanhe River Basin

## Results

### Land use change in Yanhe River Basin

Land use in 1980 and 2017 in Yanhe River Basin is shown in Table 1 and Figure 3. There is no large area of cultivated land and it is scattered in grasslands and woodlands. Grassland is located in upper and lower reaches of the basin in both 1980 and 2017, forestland is located in middle reaches in 2017. Dominant land use type is grass and cultivated land which accounts for about 88.82% of the whole area in 1980, grass and forest land takes up about 84.33% in 2017. There are two obvious land use change from 1980 to 2017: the increase of forest land, urban use land and decrease of cultivated land, grassland and waters. Compared with land use in 1980, cultivated land area is from 3348.9km^2^ to 872.10 km^2^ and the area percent decreases from 43.02 to 12.20; Forest land area ranges from 782.4 km^2^ to 3018.52 km^2^ and the area percent increases from 10.05 to 38.77. Area percent of grassland, waters, barren, urban use land changes slightly, and the value is −0.24, −0.20, 0.32 and 3.2 respectively. Land use change demonstrates that the GGP has obtained great achievements in Yanhe River Basin.

**Table 1.** Land use in Yanhe River Basin in 1980 and 2017.

| Year | Land use | Cultivated | Forest | Grass | Urban | Waters | Barren |
| --- | --- | --- | --- | --- | --- | --- | --- |
| 1980 | Area(km<sup>2</sup>) | 3348.90 | 782.40 | 3565.20 | 12.60 | 26.90 | 49.00 |
|  | Percent(%) | 43.02 | 10.05 | 45.80 | 0.16 | 0.35 | 0.63 |
| 2017 | Area(km<sup>2</sup>) | 872.10 | 3018.52 | 3547.09 | 261.84 | 11.42 | 74.03 |
|  | Percent(%) | 11.20 | 38.77 | 45.56 | 3.36 | 0.15 | 0.95 |
| change | Area(km<sup>2</sup>) | -2476.80 | 2236.12 | -18.11 | 249.24 | -15.48 | 25.03 |
|  | Percent(%) | -31.82 | 28.72 | -0.24 | 3.20 | -0.20 | 0.32 |

**Figure 3.**
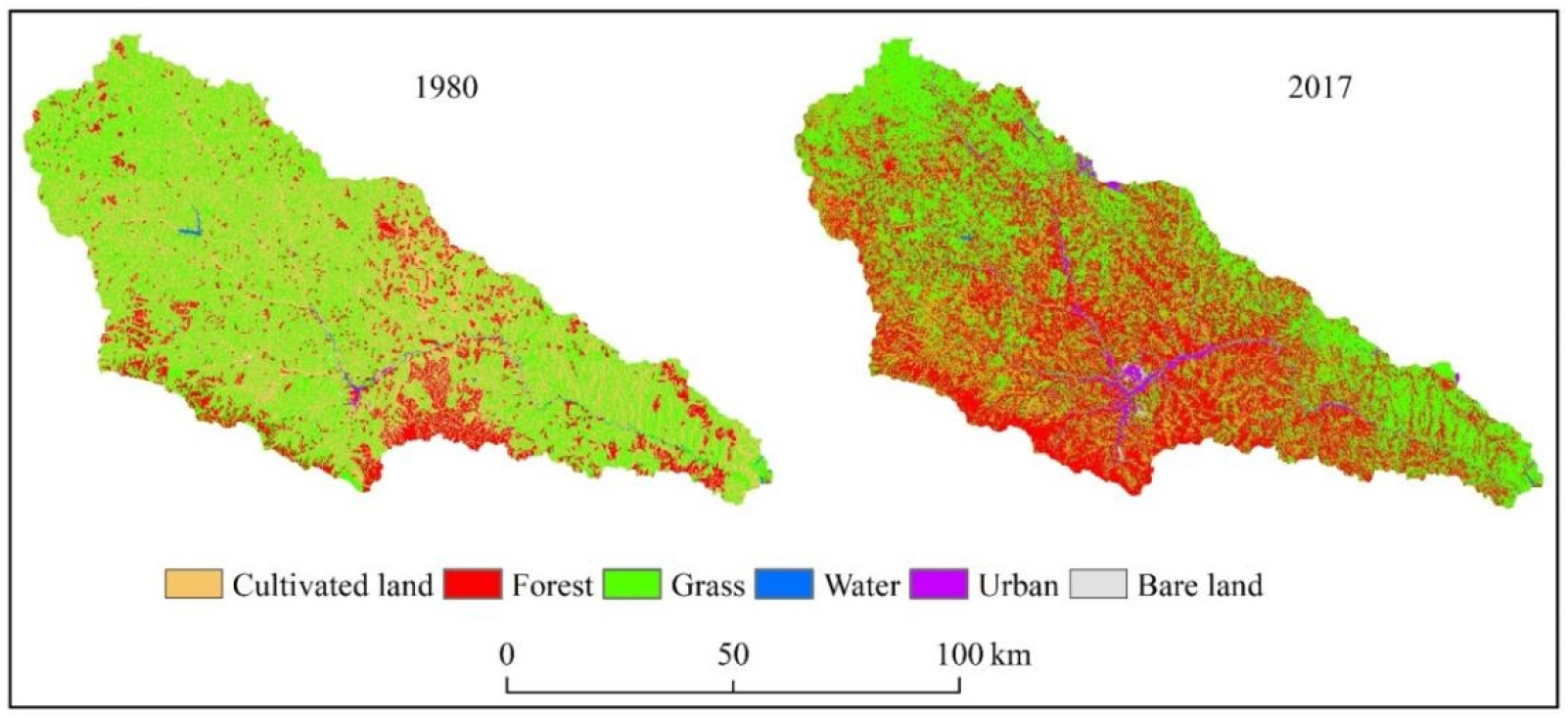
Land use map of Yanhe River Basin in different year

Diagonal value in land use conversion matrix (Table 2) is the unchanged area of every land use type. 5132.30 km^2^ of land use type has changed during the period from 1980 to 2017, 2937.70 km^2^ of cultivated land has changed into other land use types and the area proportion is 57.24%; 1198.30 km^2^ has converted into forest and 1586.44 km^2^ into grassland. Grassland is the second land use conversion type (1843.56 km^2^) which is mainly converted into forest (1337.85 km^2^), cultivated land (388.01 km^2^) and urban use land (103.85 km^2^). The transfer out land use type is cultivated land and grass land which the transfer proportion takes up as high as 93.16% of the whole basin.

**Table 2.**
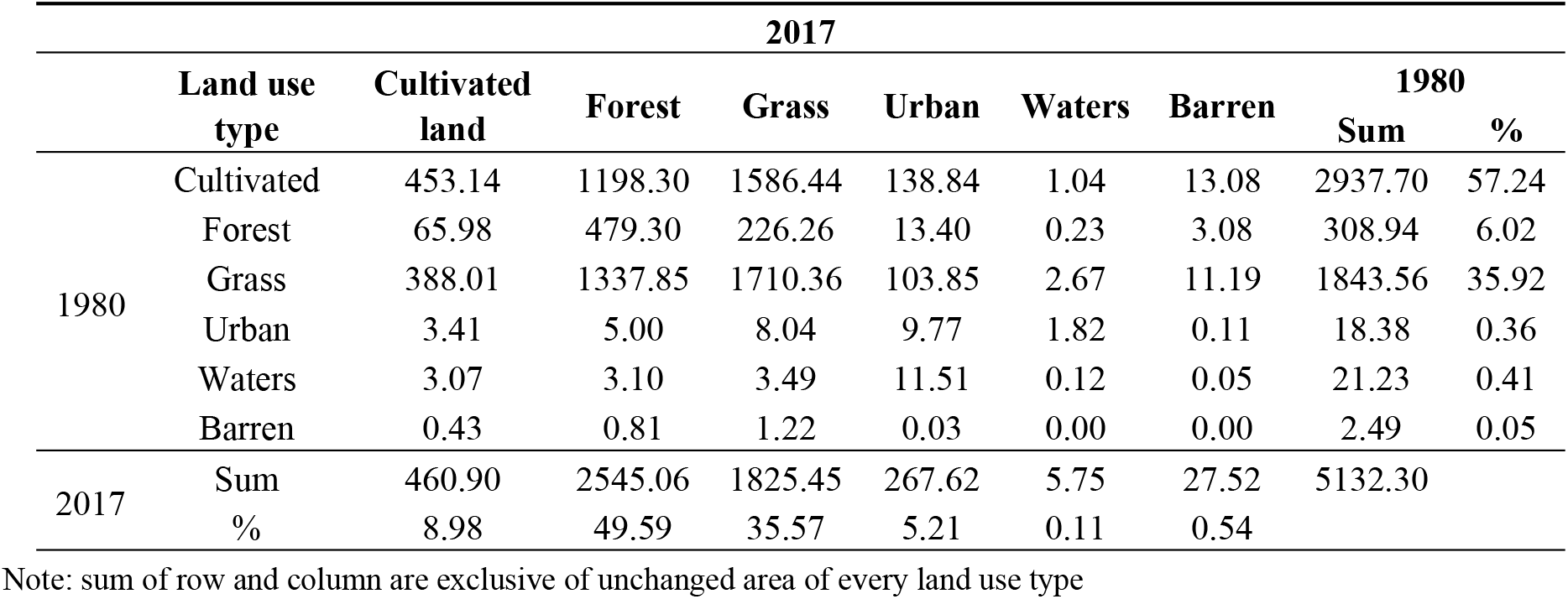
Transition matrix of land use conversion (km^2^) from 1980 to 2017.

2545.06 km^2^ of forest is developed from other land use types since 1980 and the most conversion is from grass and cultivated land; in the area transferred from other land use types, the proportion of forest accounts for 49.59%. 1825.45 km^2^ grass land is from other land use types and 1198.30 km^2^ is from cultivated land; the grassland transferred proportion is 35.57% of the whole basin. Though cultivated land mostly transferred into forest and grass land, there are 65.98 km^2^ and 388.01 km^2^ of cultivated land is from forest and grass respectively.

### Calibration and validation of the SWAT model

Sensitivity analysis indicates that CN2 (SCS streamflow curve number for moisture condition 2), CANMX (Maximum canopy storage), ALPHA_BNK (Baseflow alpha factor for bank storage), SOL_AWC (Available water capacity of the soil layer), SOL_K (Saturated hydraulic conductivity of the soil layer), SOL_BD (Moist bulk density of the soil layer), ESCO (Soil evaporation compensation factor), and REVAPMN (Threshold depth of water in the shallow aquifer for “revap” to occur) are more sensitive than other parameters for streamflow simulation and their optimal values for SWAT model are shown in Table 3.

**Table 3.** The initial ranges and final values of sensitivity parameters.

| Parameters | Rank | t-Stat | Calibrated method | P-Value | Initial range | Optimal value |
| --- | --- | --- | --- | --- | --- | --- |
| CN2.mgt | 1 | -33.69 | R | 0.00 | (-1,1) | -0.74 |
| CANMX.hru | 2 | 6.02 | V | 0.00 | (0,100) | 0.04 |
| ALPHA_BNK_rte | 3 | -4.28 | V | 0.00 | (0,1) | 0.11 |
| SOL_AWC.sol | 4 | 3.81 | R | 0.00 | (-1,1) | -0.76 |
| SOL_K.sol | 5 | -3.69 | R | 0.00 | (-1,1) | 0.13 |
| SOL BD.sol | 6 | -2.95 | R | 0.00 | (-1,1) | 0.16 |
| ESCO.hru | 7 | -2.35 | V | 0.02 | (0,1) | 0.37 |
| REVAPMN.gw | 8 | 1.67 | V | 0.09 | (0,500) | 127.83 |

Note: R means existing parameter value is multiplied by (1+ a given value), V means the default parameter is replaced by a given value.

SWAT-CUP is used to calibrate and validate the model in this study, 1980-1985 is selected as warm up period, 1986-1994 is the calibration period, 1995-1997 is the validation period. R^2^ (coefficient of determination), NSE (Nash-Sutcliffe efficiency coefficients) and percent bias (BIAS) are used to assess the accuracy of model simulation. When values of R^2^ and NSE are more than 0.5, the value of BIAS is less than or equal to ±20%, the SWAT calibration results on a monthly scale is considered to be acceptable. Figure 4 illustrates the calibration and validation results in Ganguyi hydrological station in Yanhe River Basin. R^2^, NSE, and BIAS is equal to 0.79, 0.73 and 2.20% in calibration period; R^2^, NSE, and BIAS is equal to 0.71, 0.69 and 15% in validation period respectively. The goodness-of-fit statistics indicates a reasonable agreement between observed and simulated streamflow. However, there exists higher deviation in July in 1989 and 1996 because the precipitation on 16th in July in 1989 is 26.7mm and the precipitation on 12th in July in 1996 is 91.9 mm which accounts for 49.7 percent and 98.9 percent in July of the two years which maybe the reason for comparatively higher deviation between observed and simulated streamflow. The SWAT model cannot accurately simulate rainstorm processes, heavy rain in 1989 and 1996 leads to the poor simulation of the two years and reduces the whole accuracy of the simulation and validation period.

**Figure 4.**
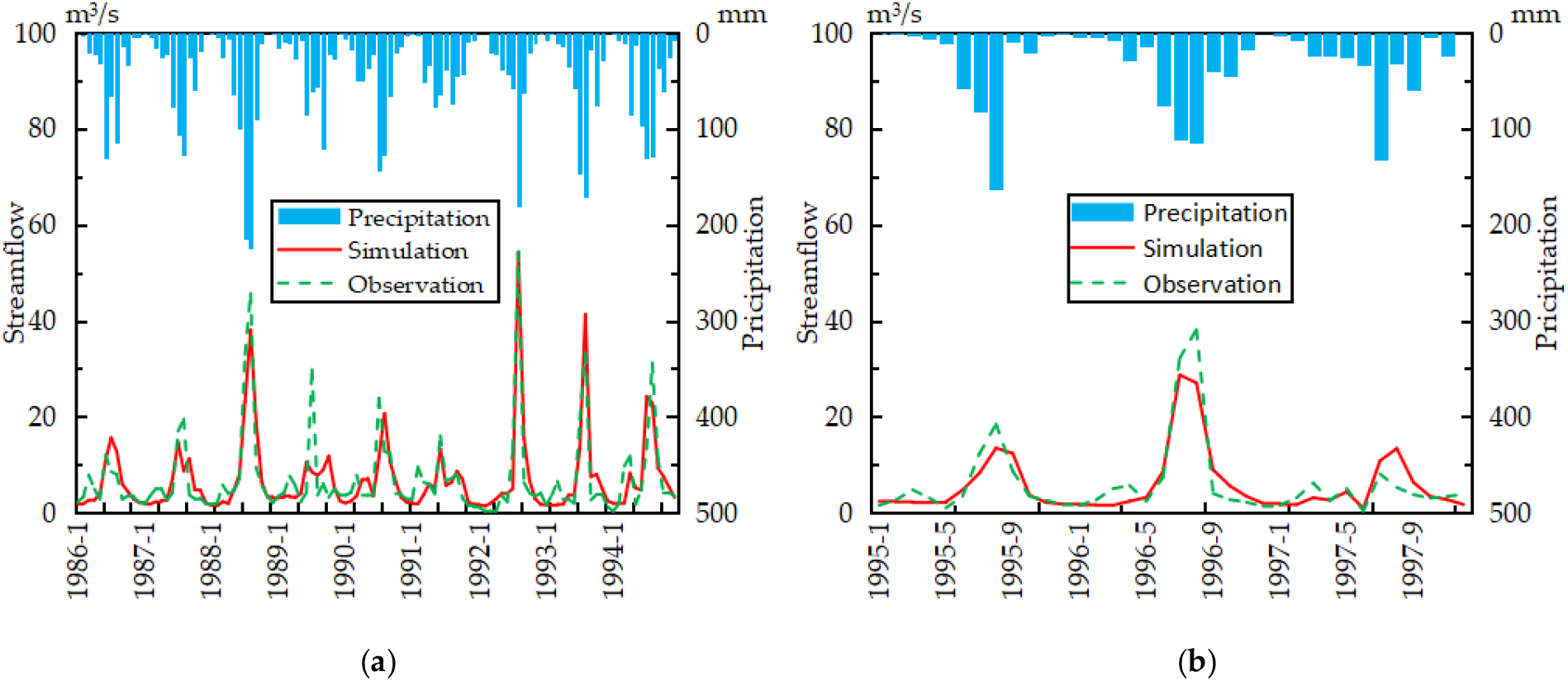
Time series plot between observed and simulated streamflow at monthly time scale

### Spatial-temporal change of blue and green water

#### Temporal change of blue and green water

##### Interannual variation of blue and green water

Temporal change characteristics of blue and green water in different vegetation cover conditions play an important role for GGP policy. The annual blue and green water simulated by SWAT model in two scenarios for Yanhe River Basin are shown in Figure5a, Figure5b. The annual average blue water resources is 13.71 billion m^3^, which accounts for 33.12% of the annual average total water resources; and the annual average green water is 27.68 billion m^3^, which accounts for 66.88% of the annual average total water resources in scenario I. The annual average blue water resources is 13.1 billion m^3^ and the annual average green water resources is 27.92 billion m^3^, which accounts for 31.93% and 68.07% of the annual average total water resources respectively in scenario II. The maximum blue water appears in 2013 and the minimum in 1999 and the two years correspond to the maximum and minimum of precipitation in the study period. Blue water in 1981, 1988, 2017 are more than that in other years and in 1995, 2000, 2004 are less than that in other years which is the same as the temporal characteristics of precipitation and indicates the amount of blue water is strongly related to rainfall, and the correlation coefficient of two elements is 0.97. There is no significant relation between the amount of rainfall and green water.

Blue water increases with a rate of 0.87 million m^3^/yr and 2.2 million m^3^/yr in scenario I and scenario II respectively; the green water increase rate is 10.29 million m^3^/yr and 10.8 million m^3^/yr in scenario I and scenario II respectively. Increase rate of blue water in scenario II is about 2.52 times that in scenario I, but there is almost no difference in increase rate of green water between two different scenarios. Both blue water and green water show increase trend in two land use scenarios.

Blue water, green water and total water resources in scenario I is less than that in scenario II in 38, 14 and 34 years in the studied 38 years (Figure 5c). Average annual difference of blue, green and total water resources between scenario II and scenario I is −72.08 million m^3^, 24.34 million m^3^, −47.74 million m^3^ respectively which indicates that land use change caused by GGP leads to decrease in blue and generalized water resources whereas increase in green water resources.

**Figure 5.**
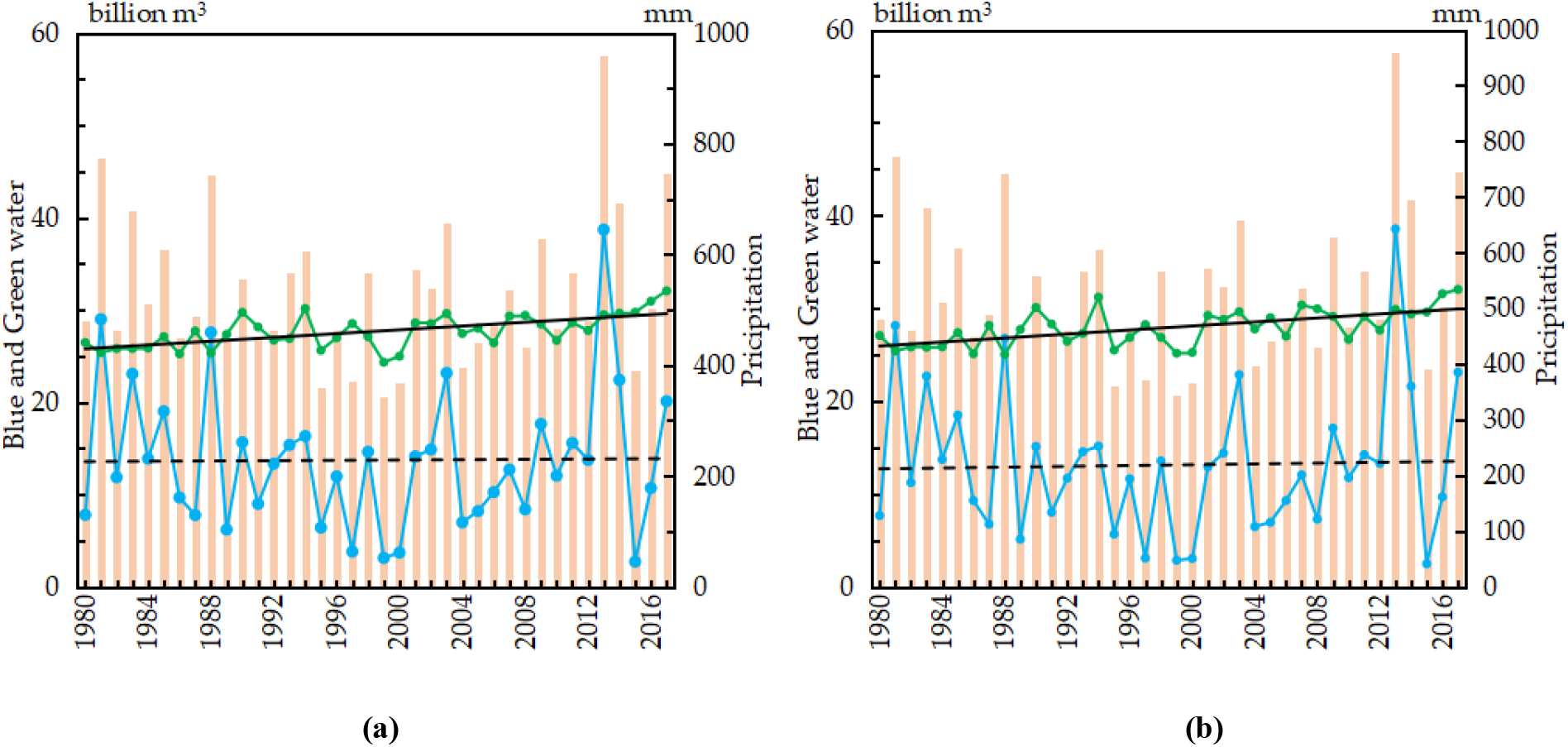
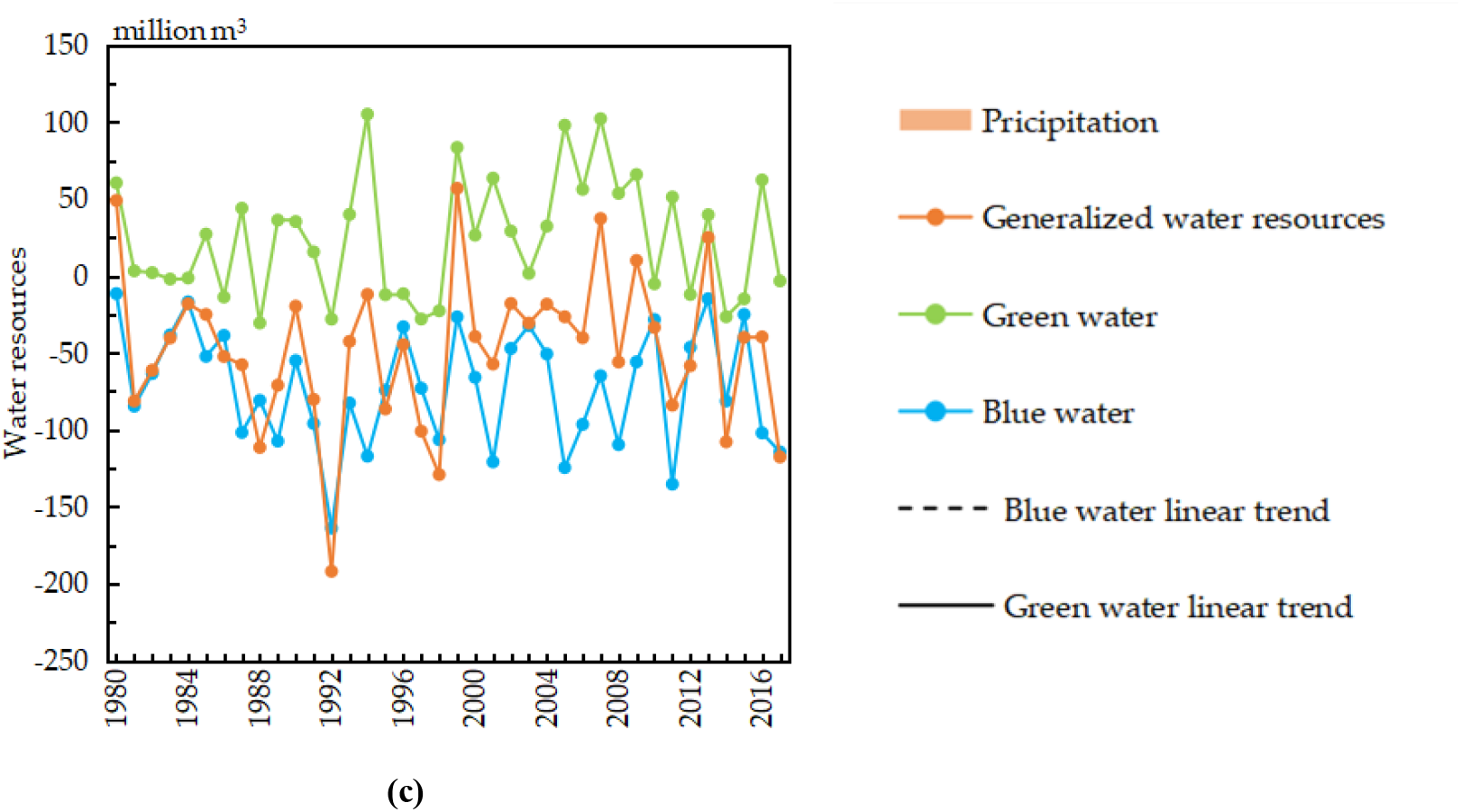
Time series change of annual water resources

##### Interannual variation of diverse hydrological components

In order to clarify impact of land use on regional water resources, it is necessary to analyze and quantify the diverse components of hydrological elements within the study area. It includes SURQ, LATQ, GWQ, ET (green water flow), SW(green water storage) obtaining from the well calibrated SWAT model. Figure 6 demonstrates that annual SURQ, component of blue water in scenario II is smaller than that in scenario I in all studied years except in 2017; the annual average difference between scenario IIand scenario I is 17.56mm and the maximum difference 45.61mm appears in 2013. Change in SURQ is consistent with the fact that surface runoff decrease in the Loess Plateau since the implementation of GGP. LATQ in scenario II is larger than that in scenario I in 2000 and 2015 but lesser in other years; change range of LATQ is the smallest in the five variables, the annual average difference between scenario II and scenario I is 2.5 mm. GWQ in scenario II is larger than that of scenario I in 1992, 1995, 2000, 2005, 2008 and 2015 and smaller in other years, the annual average difference between scenario II and scenario I is 6.57 mm.

**Figure 6.**
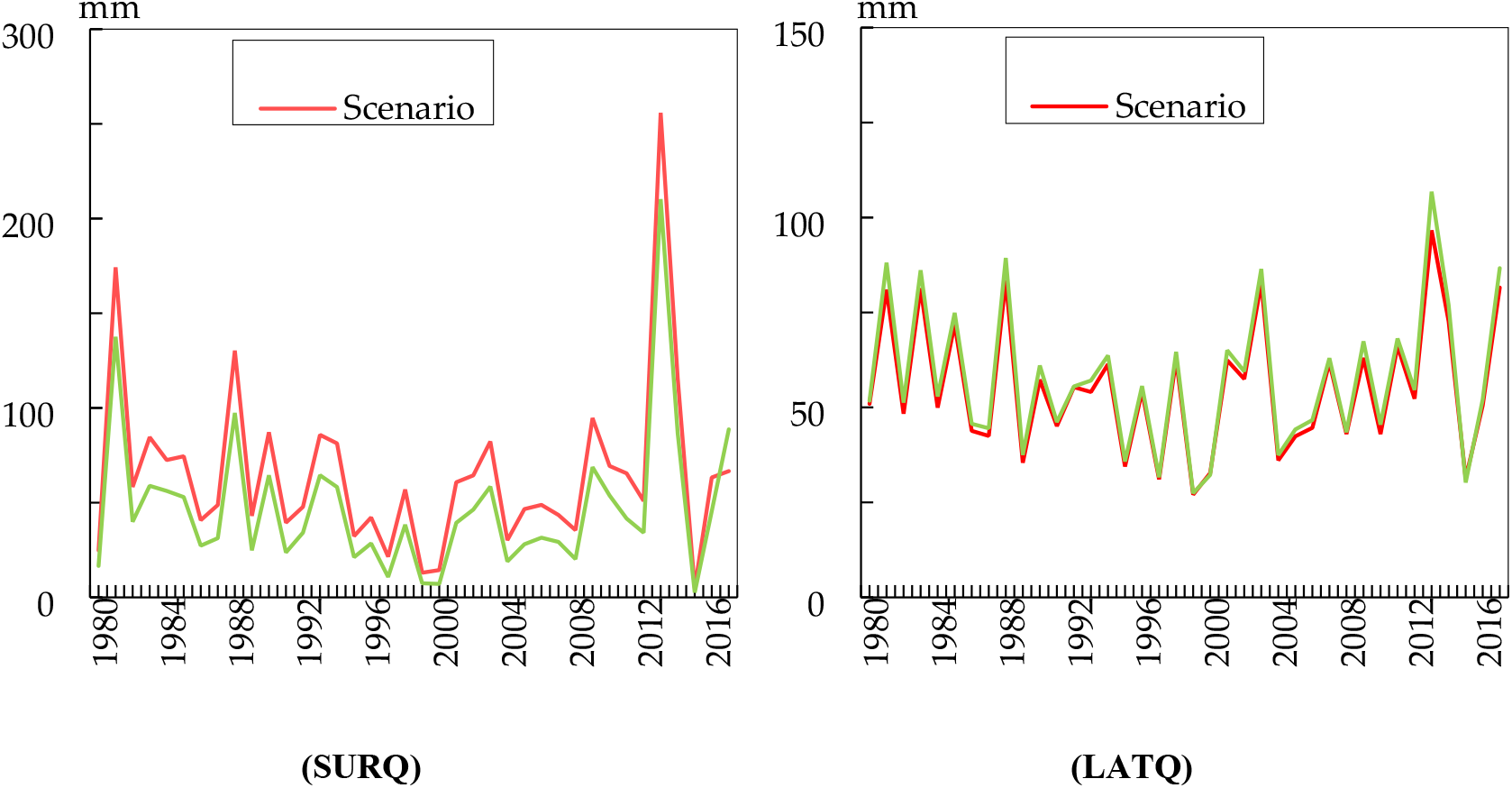
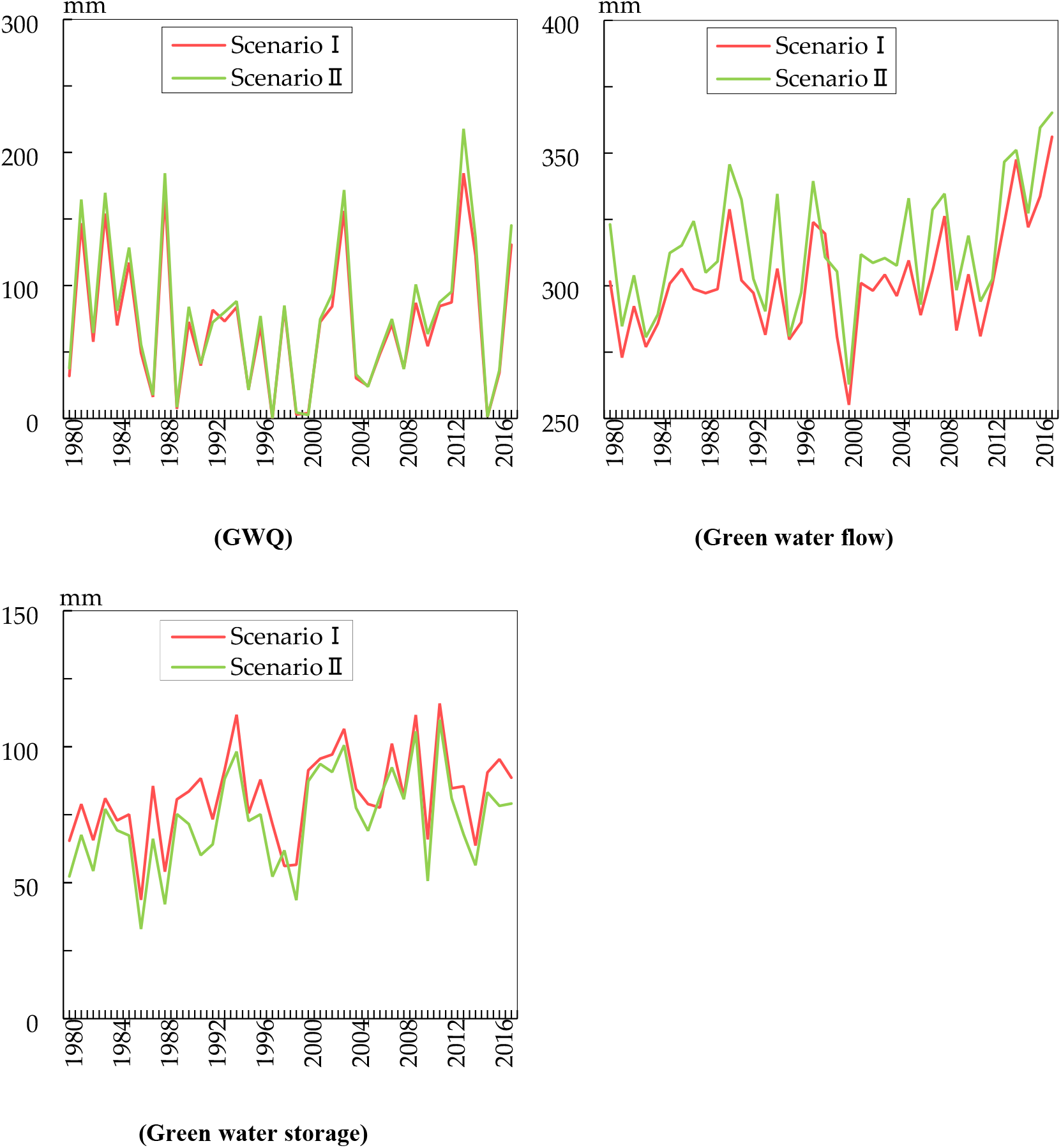
Temporal change of hydrological elements

Annual green water flow, component of green water in scenario II is larger than that of scenario I in all studied years except in 2000, whereas green water storage in scenario II is less than that in scenario I in all the studied years (Figure 6). The mean annual green water flow shows a variation from 255.15 mm to 356.12 mm in scenario I and from 262.81 mm to 365.19 mm in scenario II; The mean annual green water storage shows a variation from 43.68 mm to 115.92 mm in scenario I and from 32.96 mm to 109.92 mm in scenario II throughout the study period; green water flow in scenario I is smaller than that in scenario II except in 1998, whereas green water storage in scenario I is more than that in scenario IIexcept in 1998. The difference of green water flow between two land use scenarios is comparatively bigger in comparison to green water storage. The annual average difference of green water flow and green water storage between scenario II and scenario I are 12.28 mm, −8.9 mm respectively. GGP reduces SURQ, GWQ and green water storage but increases LATQ and green water flow.

#### Spatial distribution change of blue and green water

Blue water, green water storage, and green water flow show considerable spatial variation among sub-basins in two land use scenarios (Figure 7). Water resource values are divided into 5 ranks according to natural breaks method in ArcGIS software in scenario I, whereas the classification boundary in scenario II is just the same as that in scenario I in order to compare the spatial variations in two scenarios.

**Figure 7.**
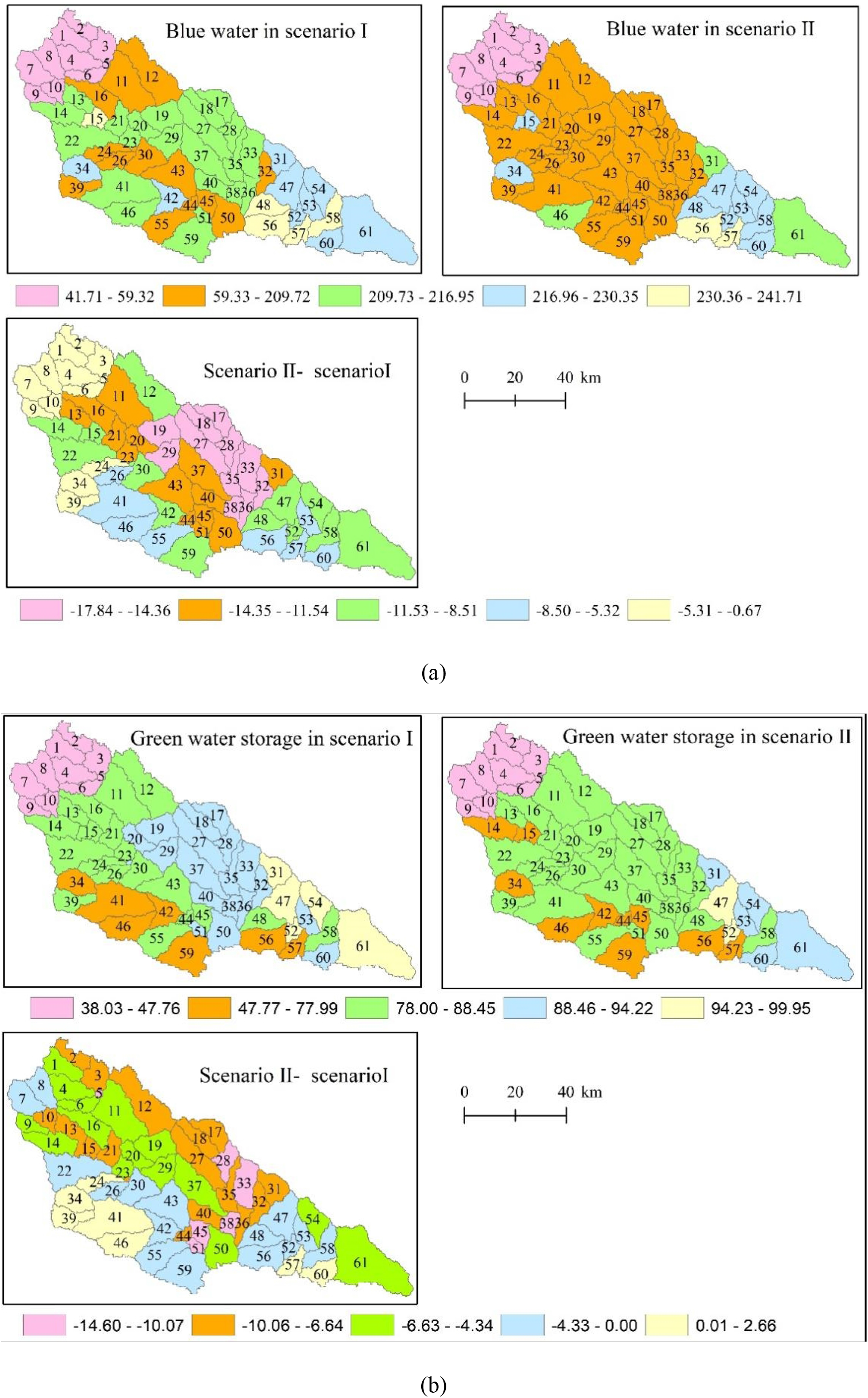
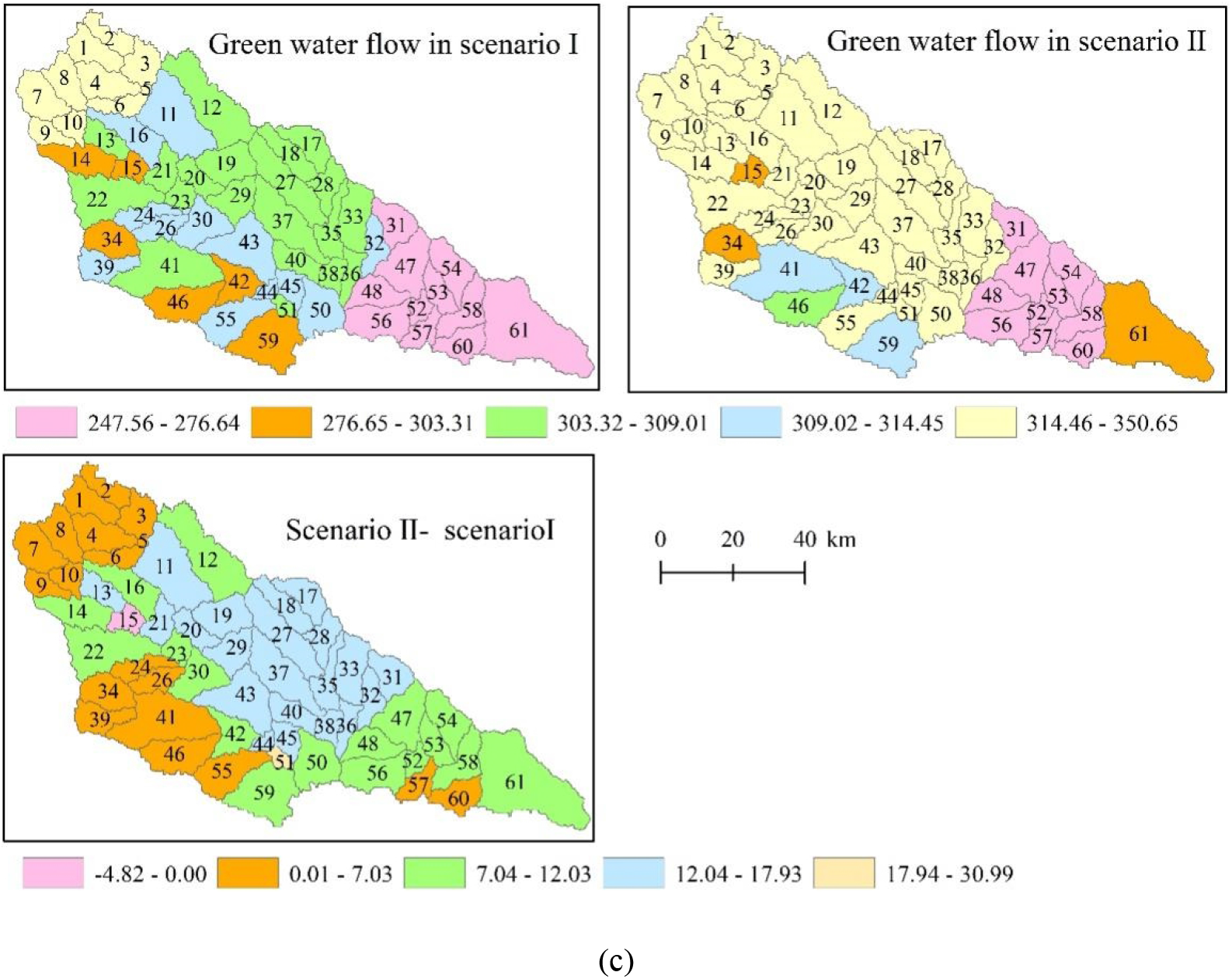
Spatial distribution of blue and green water

The spatial distribution of mean annual blue water less than 59.32 mm is exactly the same in two scenarios and the ten minimum change regions are distributed in the upper area of the basin (Figure 7a). It can be observed that blue water in second rank jumps from 59.33 mm to 209.72 mm, 14 sub-basins in scenario I and 36 sub-basins in scenario II are in the range. 37 sub-basins with average annual blue water ranging from 207.93 mm to 241.71 mm while only 15 sub-basins are in the interval in scenario II. Figure 7a presents that blue water in scenario II is less than that in scenario I in all sub-basins which demonstrates that blue water decrease in the whole basin with the implementation of GGP. The minimum change occurs in the upper part and the maximum change happens in middle part of the basin.

Figure 7b presents the green water storage across sub-basins for two scenarios. It indicates that the green water storage less than 47.76 mm is both located in the upper reaches of the river basin in two land use scenarios. Area with the value between 78 mm and 88.45 mm occupies 57.44 percent of the whole basin in scenario II. Green water storage in 8 sub-basins in scenario II is larger compared with that in scenario I and the area occupies 12.89 percent of the whole basin. The maximum decrease region is located in the north and east of the basin and the increase region is located in the west of the basin.

The maximum green water flow locates in upper reaches in scenario I but upper and middle reaches in scenario II (Figure 7c). Data in 42 sub-basins are more than 314.46 mm in scenario II and the area accounts for 63.21 percent of the whole basin; green water flow less than 276.64mm accounts for 14.75 percent in scenario II and 21.5 percent in scenario I, it indicates that there is comparatively little variability in areas of the minimum data range between two land use scenarios. Green water in 1 sub-basin decrease and the area occupies 0.79 percent of the basin, therefore, it can be regarded as that the green water flow raise in the whole basin because of GGP.

Land use conversion map from 1980 to 2017 and sub watershed in SWAT model are superimposed on one map (Figure 8) in order to compare land use conversion with spatial change of water resources. We can see that spatial change of blue water, green water storage, and green water flow (Figure 7) have relation with land use conversion to a certain extent. Area with the least reduction of blue water is consistent with area which land use unchanged, such as sub-basin from one to ten; whereas area with the largest reduction of blue water is consistent with area which land use changes significantly, such as sub-basin 17, 18, 19, 27, 28, 29, 32, 33, 35, 36, 38 where a large number of cultivated land are converted into grass and forest. The second reduction area such as sub-basin 11,13, 16, 20, 21, 23 and so on is also the area that many cultivated land are converted into grass and forest. The relation between spatial change of green water flow and land use conversion is relatively the same as between blue water and land use conversion. There is no significant relation between land use conversion and spatial change of green water storage.

**Figure 8.**
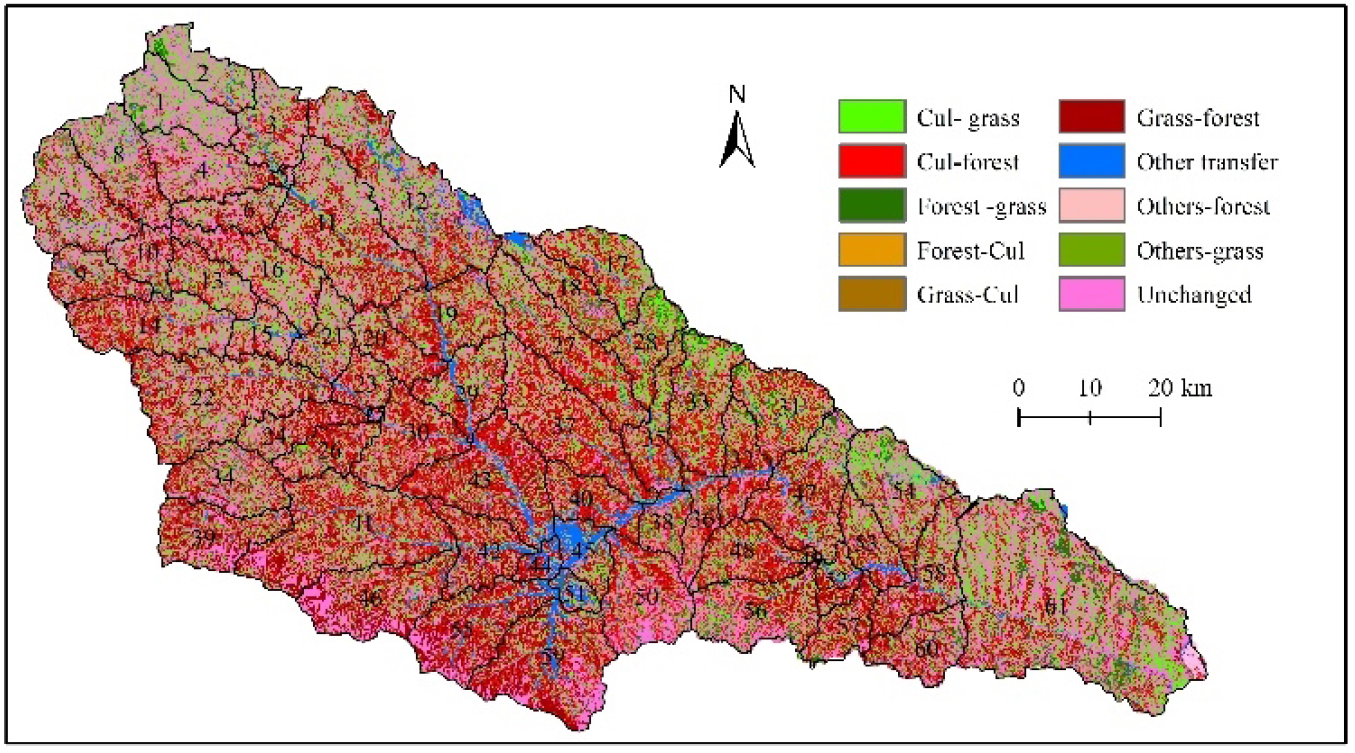
Land use conversion between 1980 and 2017

### WF

#### Agricultural WF

This paper calculates crops WF during the period of 1994-2017 because of the crop output inaccessibility before 1994 (Figure 9). Agricultural WF in Yanhe River Basin shows a rapid downward trend, with the highest value of 250 million m^3^ in 1997 and the lowest value of 136 million m^3^ in 2003. The average annual WF of agricultural products is about 160 million m^3^, with an annual decrease of about 4.07 million m^3^/yr.

**Figure 9.**
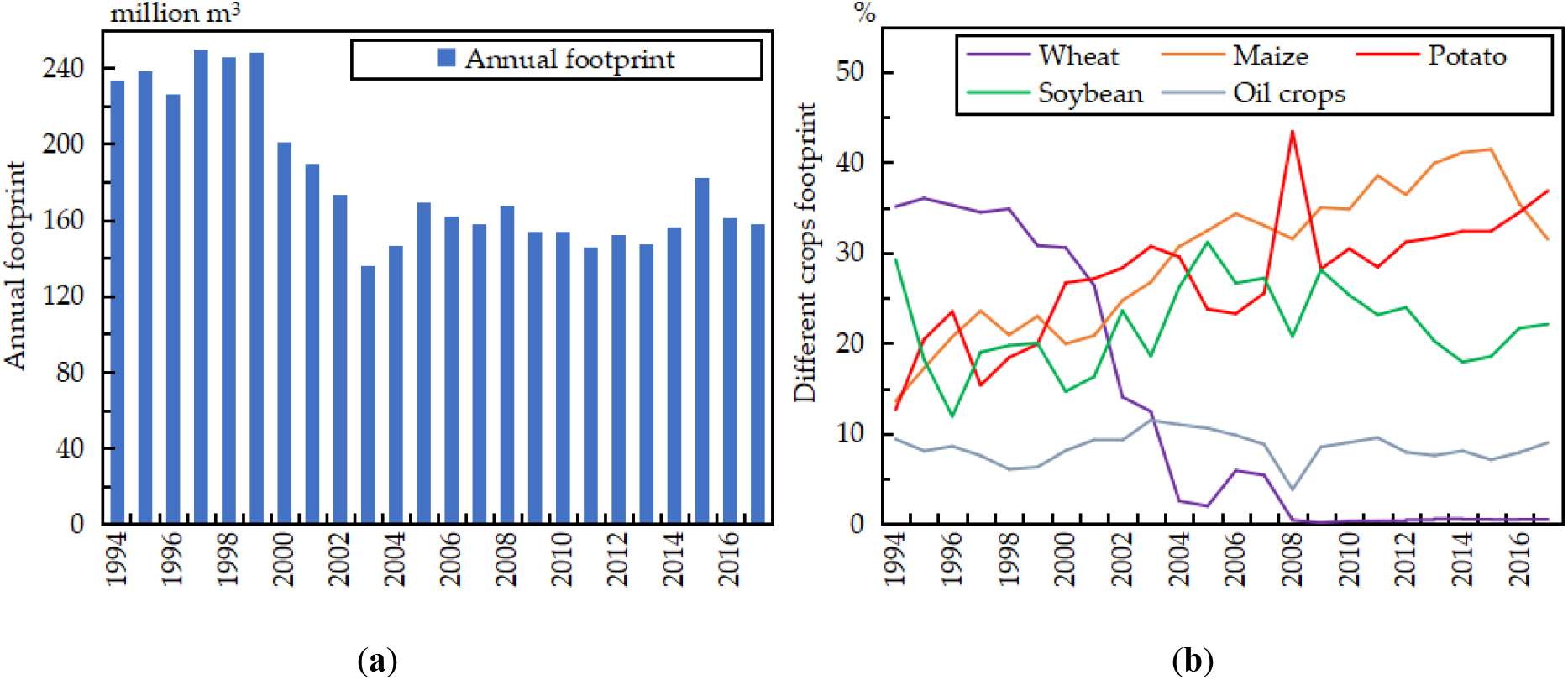
Agricultural footprint

WF of each crops in Yanhe River Basin has changed greatly during the study period. Wheat is no longer planted in Baota District after 2005 and Ansai County after 2008, and the yield in Yanchang County has decreased from 16238 tons in 1994 to 353 tons in 2017. Therefore, wheat WF has decreased from 81.97 million m^3^ in 1994 to 0.84 million m^3^ in 2017, and the percent is from 35.1 to 0.52. WF proportion of corn and potato both shows an obvious upward trend, and corn WF increases from 31.75 million m^3^ to 50.58 million m^3^, the proportion increases from 13.6 percent to 31.5 percent. The potato WF increases from 29.52 million m^3^ to 59.13 million m^3^, the proportion increases from 12.65 percent to 36.85 percent. Soybean and oil crops WF fluctuate in different years, but the overall change trend is not obvious in the study period.

#### Ecological WF

Vegetation WF is calculated according to equation (7) and (8) in ArcGIS by vegetation type, soil texture and potential evaporation, and results are demonstrated in Table 4. It presents that forest WF is larger in May, June and July because the actual evapotranspiration of the three months is more than that in other months, and the minimum value appears in October because trees begin to wither and fall leaves, water demand is the smallest in the growth stage. Total forest WF increases from 5.22 billion m^3^ to 19.58 billion m^3^ during 1980-2017. After 2000, forest WF begins to increase rapidly and the maximum value appear in 2017.

**Table 4.** WF of forest from April to October in different years(billion m^3^)

| Month | 1980 | 1995 | 2000 | 2005 | 2010 | 2017 |
| --- | --- | --- | --- | --- | --- | --- |
| 4 | 0.74 | 0.81 | 0.91 | 1.32 | 2.52 | 3.35 |
| 5 | 0.85 | 1.04 | 1.17 | 1.40 | 3.02 | 4.03 |
| 6 | 0.95 | 0.96 | 0.89 | 1.52 | 3.31 | 3.74 |
| 7 | 0.86 | 0.85 | 0.99 | 1.30 | 2.71 | 3.56 |
| 8 | 0.74 | 0.56 | 0.67 | 0.93 | 2.05 | 2.31 |
| 9 | 0.62 | 0.43 | 0.50 | 0.69 | 1.48 | 1.73 |
| 10 | 0.46 | 0.27 | 0.24 | 0.45 | 1.05 | 0.87 |
| Summary | 5.22 | 4.92 | 5.37 | 7.6 | 16.14 | 19.58 |

Grass WF is larger in May and June but small in October in grass growth stage which is just the same as the monthly characteristic of forest. Table 1 has demonstrated that grassland area decreased from 3565.2 km^2^ to 3547.09 km^2^ which indicates that GGP has comparatively little effect on grassland so grass WF has no obvious change and decreases slightly from 18.71 billion m^3^ to 15.44 billion m^3^ (Table 5).

**Table 5.** WF of grassland from April to October in different years(billion m^3^)

| Month | 1980 | 1995 | 2000 | 2005 | 2010 | 2017 |
| --- | --- | --- | --- | --- | --- | --- |
| 4 | 2.66 | 2.33 | 3.04 | 3.33 | 2.43 | 2.64 |
| 5 | 3.04 | 3.15 | 3.86 | 3.55 | 2.92 | 3.18 |
| 6 | 3.41 | 3.41 | 2.97 | 3.85 | 3.21 | 2.95 |
| 7 | 3.09 | 3.20 | 3.23 | 3.15 | 2.63 | 2.81 |
| 8 | 2.66 | 2.80 | 2.21 | 2.31 | 1.98 | 1.82 |
| 9 | 2.21 | 1.98 | 1.63 | 1.66 | 1.43 | 1.37 |
| 10 | 1.64 | 1.38 | 0.73 | 1.04 | 1.02 | 0.68 |
| Summary | 18.71 | 18.26 | 17.67 | 18.90 | 15.61 | 15.44 |

#### Agricultural Virual water flow

Figure 10 indicates that from 1994 to 2017, grain consumption WF in Yanhe River Basin ranged from 1.01 billion m^3^ to 3.24 billion m^3^. In 1994-1997, 2002-2003, 2012 and 2014-2017, the virtual water was in outflow state, while in other years, it was in inflow state. From 1994 to 2017, the average annual inflow of virtual water was 8 million m^3^. Therefore, in general, the virtual water quantity of grain in Yanhe River Basin is imported from outside which can alleviate the pressure of water resources in the basin to a certain extent.

**Figure 10.**
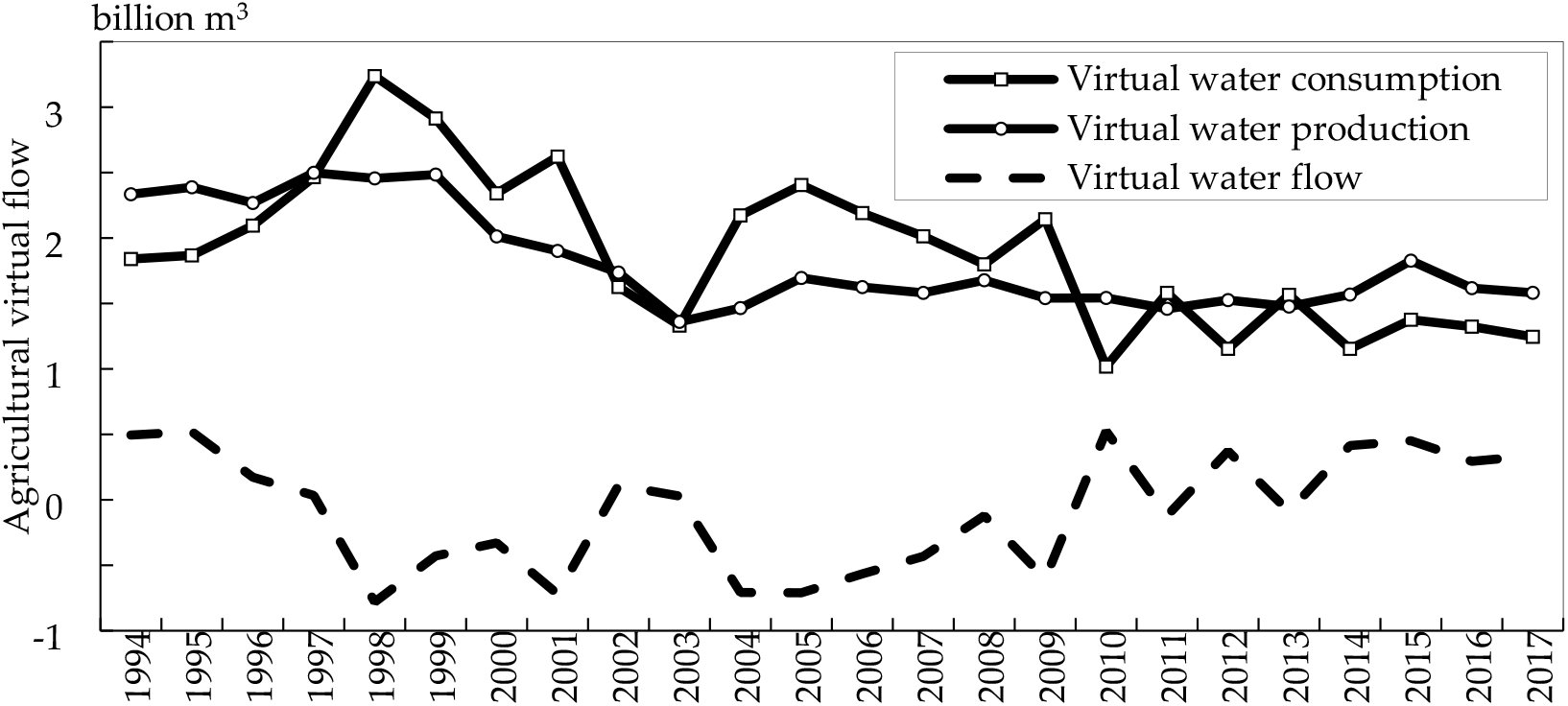
Agricultural virtual flow during 1994-2017

#### EWSI

EWSI is calculated in 1995, 2000, 2005, 2010 and 2017 and the results are shown in Table 6. Total WF shows increase trend due to the significant increase of forest WF though agricultural and grass WF show decrease trend during the study years. However, ecological water stress index shows no obvious temporal change because change trend of generalized water resources is the same as that of regional total WF. EWSI is less than 1 in the 5 studied years in both two land use scenarios from the point view of generalized water resources. ESWI in scenario II is bigger than that in scenario I in the 5 years and it can demonstrate that GGP has slightly increased regional water stress because difference of ESWI is about 0.01 between two land use scenarios.

**Table 6.** Ecological water stress index in two land use scenarios in Yanhe River Basin.

| Year | WF<br>(billion m<sup>3</sup>) | Scenario I |  | Scenario II |  | Scenario II - I |
| --- | --- | --- | --- | --- | --- | --- |
|  |  | Generalized<br>water<br>resources<br>(billion m<sup>3</sup>) | ESWI | Generalized<br>water<br>resources<br>(billion m<sup>3</sup>) | ESWI | ESWI |
| 1995 | 25.2 | 35.45 | 0.711 | 34.50 | 0.730 | 0.02 |
| 2000 | 26.32 | 31.75 | 0.829 | 31.31 | 0.841 | 0.01 |
| 2005 | 31.26 | 40.08 | 0.780 | 39.79 | 0.786 | 0.01 |
| 2010 | 36.12 | 42.88 | 0.842 | 42.51 | 0.850 | 0.01 |
| 2017 | 35.78 | 46.10 | 0.776 | 45.66 | 0.784 | 0.01 |

## Discussion

This paper investigates the spatial-temporal impact of GGP on regional water resources and ecological water stress in Yanhe River Basin in 1980 and 2017. In this study, accuracy of SWAT simulation is one of the important key factors because many results such as blue water, green water, and hydrological elements (green water flow, green water storage, SURQ, GWQ and LATQ) are all output of the model. R^2^ and NSE in this paper are within the accuracy requirement of model establishment, and the accuracy is close to the results of other researches in this river basin [53–55]. During the SWAT simulation, land use is classified into 6 types which cannot present the land use type actually and the land use module should be improved. 4 meteorological stations such as Yanchang, Ansai, Zhidan and Yanan are input in SWAT and precipitation data are also from the meteorological stations, more meteorological and precipitation data may enhance simulation accuracy. Reservoir such as Wangyao information can be input in the model so as to consider human impact and improving DEM quality can also improve simulation accuracy [56].

Only 5 crops such as wheat, summer maize, potato, soybean and oil crops are considered while other agricultural WF are not included because of lower yield in Yanhe River Basin, therefore, agricultural WF is less than actual total agricultural WF in the region. The most important varieties of trees in the the Loess Plateau is robin pseudoacacia, platycladus orientalis, pinus tabulaeformis and so on. Different experiments on forest in the Loess Plateau show that ecological water demand coefficient of arbor forest is 0.757, and that of shrub forest is 0.612 [57]; Water demand coefficient of forest land is 0.76, that of shrub forest is 0.61, and that of sparse forest land is 0.48 [58–59]. There are no precise and uniform water demand coefficient of forest and grass in the Loess Plateau because there are few experimental studies on it. This paper cites the existing study results and does not distinguish forest types because of lacking experiment and image interpretation accuracy. The above factors will all influence the ecological WF and EWSI.

Results in this study show that blue water resources decrease and green water increase obviously, but generalized water resources decrease slightly; SURQ, SW decrease but ET increase significantly in the whole basin with the achievements of GGP; these results are similar with Yang [53] and Yang [60] in Yanhe River Basin. Large amount of cultivated lands transform into forest lands which has reduced the slope surface runoff and ultimately, improved green water storage, which is important for restoring vegetation within this region [61]. The reason for blue water decrease and green water increase maybe shift from the former to the latter. With the increase of vegetation coverage in Yanhe River Basin, the accumulation of bark debris and leaf litter can significantly increase soil surface roughness, slow down runoff rate, increase infiltration and intensify evaporation [62]. Figure 11 shows that annual maximum leaf area index (LAI) increases from 0.9 to 1.75 during 2000-2014. Annual maximum LAI show increase trend since 2000 and larger LAI can increase the interception capacity and rain loss, water evaporates before infiltration, increases green water flow and reduces the surface runoff.

**Figure 11.**
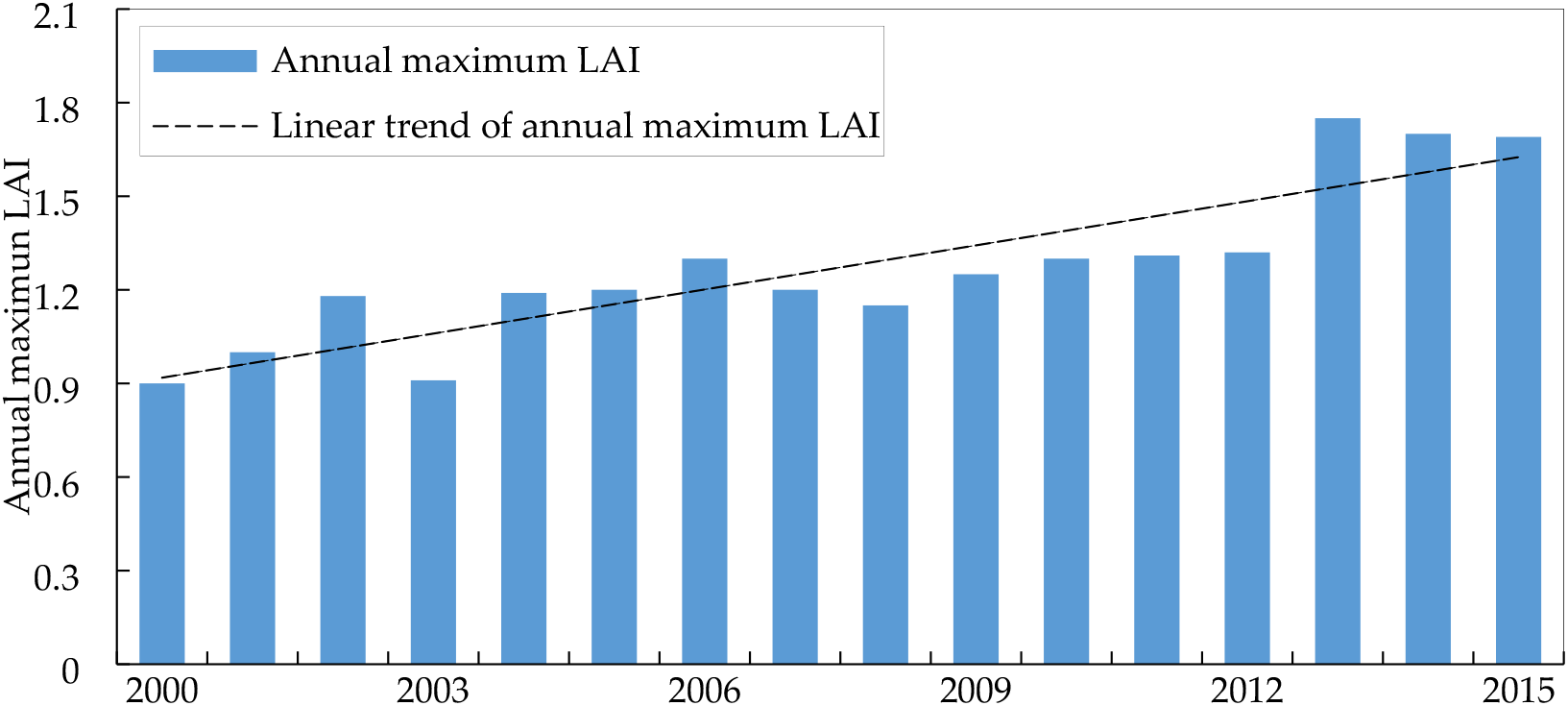
Change trend of annual maximum LAI for Yanhe River Basin during 2000-2015

Water demand of increasing vegetation in semi-arid climate of the Loess Plateau have reduced soil water content of 3-5 m to almost withering humidity, and further exacerbated soil drying [63], ET in growing season limited vegetation growth [64]. Water resources can’t satisfy water requirement in the process of forest and grass growth which leads to slow growth of trees and low yield of forest because precipitation in this region has not significantly increased during the study period but evaporation increased. The Chinese government plans to invest another US$9.5 billion in GGP on the Loess Plateau by 2050 [65]. Results in this study present that although this programme has contributed to increase forest land area and green water resources, this has been at the cost of detectable reduction in blue water resources especially in river runoff. Studies have indicated that current vegetation productivity in the Loess Plateau is already close to Net Primary Productivity (NPP) and total water resources will inevitably reduce for human use to less than the amount required if vegetation coverage continue to increase [22]. Therefore, future GGP should consider the water stress and other negative impact.

## Conclusions

This paper evaluates the impact of GGP on generalized water, blue water, green water, hydrological elements and regional water stress in Yanhe River Basin based on SWAT model and WF concept. The conclusions can be summarized as follows:

1. From 1980 to 2017, the cultivated land in the Yanhe River Basin decreased from 3348.90 km^2^ to 872.10 km^2^ and the percent is from 43.02 to 11.20, the forest land area percent increased by 27.72, urban use land area percent increased by 3.2 but other land use type areas change slightly. Vegetation coverage has been enhanced significantly since the implementation of GGP.
2. There are 38, 14 and 34 years in the study period which blue water, green water and total water resources in scenario I is less than that in scenario II. Average annual difference of blue, green and total water resources between scenario II and scenario I is −72.08 million m^3^, 24.34 million m^3^, −47.74 million m^3^ respectively which indicates that land use change caused by GGP leads to decrease in blue and generalized water resources whereas increase in green water resources.
3. Annual green water flow in scenario II is larger than that of scenario I in all studied years except in 2000, whereas green water storage in scenario II is less than that of scenario I in all the studied years. The annual average difference of green water flow and green water storage between scenario IIand scenario is 12.28 mm, −8.9 mm respectively. GGP reduces SURQ, GWQ and green water storage but increases LATQ and green water flow.
4. EWSI shows no obvious temporal change because change trend of generalized water resources is the same as that of regional total WF. It is less than 1 in 1995, 2000, 2005, 2010 and 2017 in both two land use scenarios from the point view of generalized water resources. ESWI in scenario II is bigger than that in scenario I and it presents that GGP has slightly increased regional water stress because difference of ESWI is about 0.01 between two land use scenarios.

## Acknowledgments

We thank the reviewer and editor for their insightful and constructive critiques and suggestions which helped to improve this paper.

## Author Contributions

**Conceptualization:** Yuping Han, Huiping Huang

**Methodology:** Huiping Huang, Fan Xia, Wenbin Mu

**Visualization:** Yuping Han, Fan Xia

**Funding acquisition:** Huiping Huang, Yuping Han

**Validation:** Huiping Huang, Yuping Han

**Formal analysis:** Yuping Han, Wenbin Mu

**Supervision:** Yuping Han, Wenbin Mu

**Investigation:** Huiping Huang, Fan Xia

**Writing-original draft preparation:** Yuping Han

